# Donor-derived airway ALI model for high-throughput screening of antiviral combinations with concurrent analysis of antiviral efficacy and epithelial toxicity using ciliR

**DOI:** 10.1101/2025.03.28.645652

**Authors:** Daniela Cardinale, Oriane Grant, Dani Do Hyang Lee, Robert E. Hynds, Richard Angell, Chris O’Callaghan, Mario Cortina-Borja, Elaine Thomas, Claire M. Smith

## Abstract

Respiratory RNA viral infections significantly impact quality of life and productivity, both in terms of pandemics (e.g., SARS-CoV-2, influenza A viruses) and seasonal infections such as respiratory syncytial virus (RSV). Globally, RSV causes an estimated 33.8 million cases annually in children under five, leading to 2.8–4.3 million hospital admissions and up to 199,000 deaths. Human-relevant drug screening models are key to addressing the urgent need for effective antiviral treatments for RSV, particularly in vulnerable patient groups.

Here, we present a donor-derived differentiated primary human nasal epithelial cell, 96-HTS Transwell air-liquid interface (ALI) model designed to investigate the effects of combination antiviral therapies on RSV infection in primary human ciliated airway epithelium. Additionally, we describe novel analytical tools using R (`ciliR`) to screen drug combinations by concurrently measuring efficacy and ciliary beat frequency as a sensitive marker of cell toxicity.

Our results demonstrate that the smaller 96-HTS ALI cultures retain comparable epithelial composition and ciliary function to conventional ALI culture formats. These cultures are permissible to infection with an RSV-GFP reporter virus, enabling quantitative comparison and combination treatment across multiple epithelial cultures from the same donor. We anticipate that our disease-relevant system will serve as a foundation for larger-scale experiments aimed at optimizing combination therapy for RSV and other respiratory viruses.

## Introduction

Respiratory viruses cause major morbidity and mortality worldwide. RNA viruses, including the novel coronaviruses (SARS-CoV-2) and common seasonal viruses such as respiratory syncytial virus (RSV) result in serious respiratory infections. Seasonal RSV causes major morbidity with an estimated 20,000 children under 2 years admitted to hospital each winter in the UK [1, 2]. RSV is also a significant cause of morbidity and mortality in adults over 60 years of age, causing annual epidemics in care homes during the winter season [3]. The identification of effective therapies could have a major impact on disease burden caused by RSV.

Experience from other viral infections has demonstrated that many successful therapies emerge from combining drugs with different mechanisms of action. For example, combination therapy has proved to be very successful for the treatment of hepatitis C virus (HCV) [4–6], HIV [7], herpes simplex virus (HSV) [8, 9], poliovirus [10], Ebola virus [11], Zika virus [12], and human cytomegalovirus (HCMV) [13]. Though there are compounds in development against RSV, some already in Phase 2 clinical trials, all of these are being evaluated as single therapies. The additive and/or synergistic effects of different drugs may be advantageous, improving efficacy, reducing risk of resistance emerging, with the potential of reducing drug doses required and improving therapeutic risk benefit. However, the complexities of combination clinical studies to assess compounds with different mechanisms of action in patients with an acute infection make it vital to develop appropriate higher-throughput human-relevant *in vitro* assays to measure combination effects, emergence of resistance and potential toxicity.

Small animal models have been shown to have limited utility for clinically relevant investigations into RSV therapies due to the lack of complete RSV replication [14] as well as no/limited signs and symptoms of disease. Screening in disease relevant human tissue is a key component of modern drug discovery [15]. Many respiratory viruses, including RSV [16, 17] and SARS-CoV-2 [18, 19] have been shown to target human respiratory mucosecretory and ciliated cells for infection. Ciliated cells line the respiratory tract and possess motile cilia that help to clear pathogens and debris from vulnerable cells. However, these cells are not present in standard cell line cultures, which lack the donor-specific variability seen in primary human tissue. This variability is crucial, as individual differences in cilia function and immune response can influence infection risks and drug efficacy. Therefore, using primary differentiated cell cultures derived from different human donors provides a more representative model for assessing drug efficacy, capturing the diversity in human response that may impact clinical outcomes.

Air-liquid interface (ALI) cell culture is an established method for the growth of differentiated primary human airway epithelial cells for drug screening. We have developed a human primary cell culture method that allows extensive propagation of airway basal progenitor cells [20]. This method increases the numbers of differentiated cultures that can be grown from a single biopsy and makes 96-HTS ALI culture feasible [21]. Here, we describe analysis pipelines using a 96-Transwell *in vitro* assay of differentiated human ciliated epithelial cells, as a method to test the efficacy of combination therapy using small molecule inhibitors of RSV (a schematic of the method is shown in **Figure 1**) [21].

**Figure 1:**
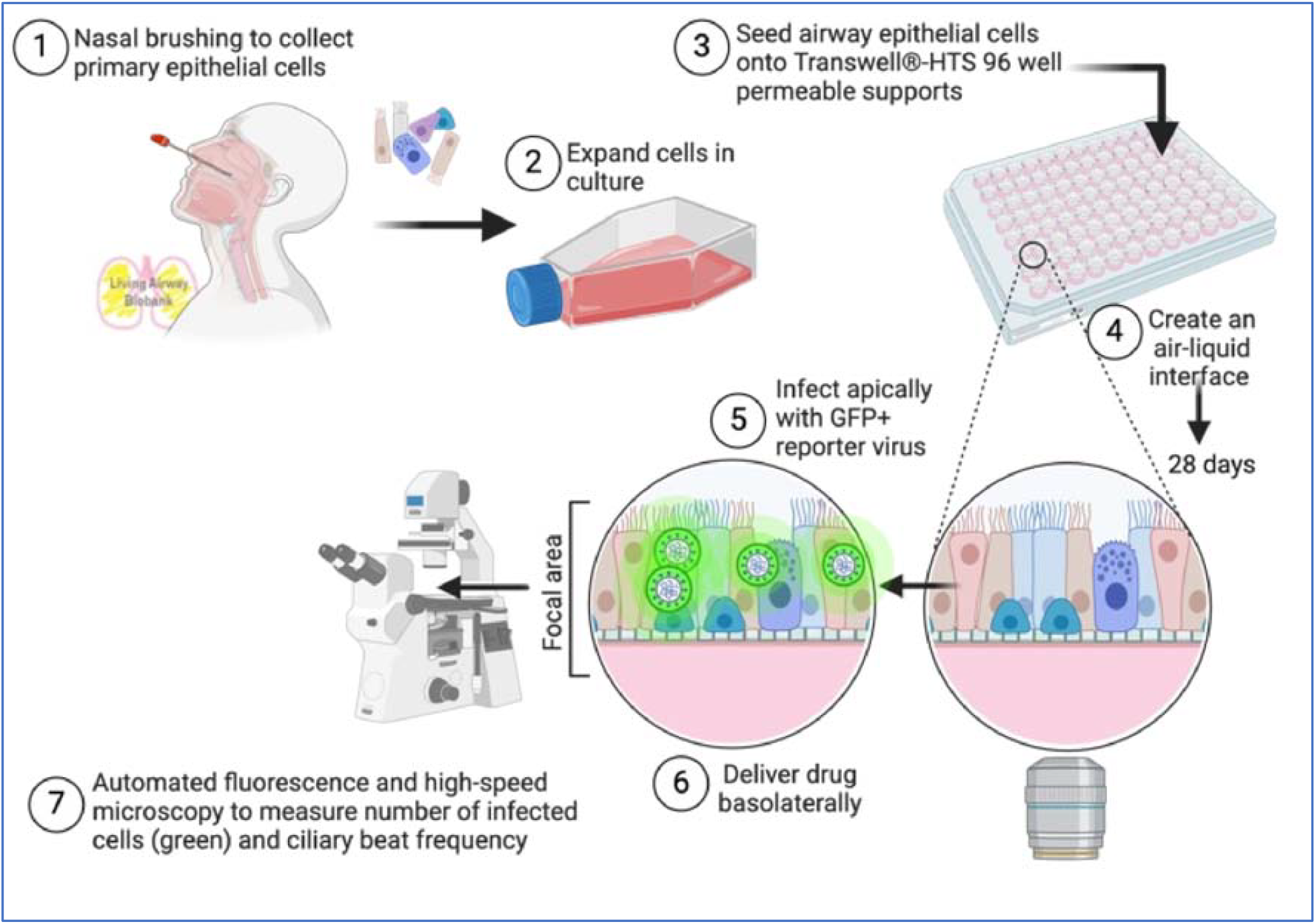
Graphical abstract of the model and method used. Image created using BioRender.com.

## Materials and methods

### Subjects

Informed written consent was obtained from all participants prior to enrolment in the study. Nasal epithelial brush biopsies were collected from healthy adult participants with ethical approval for the study provided by the UCL Living Airway Biobank (REC reference 14/NW/0128) or UCL Research Ethics (reference 4735/001).

### Culture and Differentiation of Human Airway Epithelial Cells at Air-Liquid Interface Using HTS Transwell-96 Permeable Supports

Nasal brush biopsies were received on ice in transport medium which consisted of medium 199 (Life Technologies; #22340020) containing 100 U/ml penicillin, 100 µg/ml streptomycin (Gibco; #15290026), 25 µg/ml amphotericin B and 20.5 µg/ml sodium deoxycholate (Gibco; #15290018). Cells were plated for expansion directly into co-culture with mitomycin-inactivated 3T3-J2 fibroblasts as previously described [20–22]. Briefly, basal epithelial cells were cultured at 37°C and 5% CO_2_ until confluence and then separated from feeder cells using differential trypsinization [20]. Basal cells were then seeded on collagen I-coated, semi-permeable membrane supports in submerged culture in serum-free Airway Epithelial Cell Growth Medium (Promocell, Heidelberg, Germany). For 24-well Transwells (Corning 0.4 µm pores), 1 × 10^6^ cells were seeded per membrane in 250 µl medium. For HTS Transwell-96 permeable supports (Corning; 3380; 1 µm and 0.4 µm pores, polyester membrane) 0.2 × 10^6^ cells were seeded per membrane in 75 µl medium. After 24-48 hours apical fluid was removed, and cells were fed only from the basolateral side with serum-free Airway Epithelial Cell Growth Medium (Promocell, Heidelberg, Germany). Medium was exchanged three times per week for 28 days and apical side gentle washed weekly with medium to remove mucus.

### Immunofluorescence Staining and Microscopy

The Transwell membrane was incubated in 4% (w/v) paraformaldehyde for 30 minutes at room temperature. Cells were stored at 4**°**C in PBS until the time of staining. Cells were blocked and permeabilized using a blocking buffer (3% BSA in PBS containing 0.01% Triton X) at room temperature for 1 hour, prior to overnight staining with primary antibody (in 1% BSA in PBS) at 4°C. Primary antibodies used were anti-β tubulin (Abcam; ab15568; 1:100) to detect ciliated cells and anti-MUC5AC (Invitrogen; 1:100) to detect mucin. Cells were washed 3 times in PBS for 5 minutes and secondary antibody (in 1% BSA in PBS; Molecular Probes; AlexaFluor dyes) was applied for 2 hours at room temperature. Hoechst 33258 staining solution (Sigma) was applied for 20 minutes at room temperature as a nuclear counterstain prior to imaging. For high magnification imaging, cells were mounted in 80% glycerol, 3% n-propylgallate (in PBS) mounting medium and images were obtained using an inverted LSM 710 confocal microscope (Zeiss).

### Flow Cytometric Evaluation of Cell Populations

At day 28 post ALI cell cultures were washed with PBS. Accutase (Gibco, Thermo Fisher Scientific) was added to the apical and basal sides and cultures incubated at 37 °C for 15 min. DMEM containing 10% FBS was added to the cell suspension, and then centrifuged at 400 ×g for 5 min. Cell pellets were resuspended in a FACS buffer (Phosphate Buffered Saline, 1% BSA, 0.05 mM EDTA) containing Fc receptor blocker (BioLegend) for 5 min on ice. Cells were then centrifuged again at 3400 × g for 3 min and resuspended on ice in FACS buffer containing antibodies for 20 min against: the basal cell markers integrin α6 (CD49f-PE; BioLegend) and anti-nerve growth factor receptor (NGFR/CD271; BV421-conjugated; BD Biosciences) and cell markers CD66c (for secretory epithelial cells) and Promonin-1 (Prom1/CD133) as a marker for ciliated cells [23]. After centrifugation at 3400 × g for 3 min the cell pellets were resuspended in PBS containing viability dye (BioLegend) for 10 min on ice. Viability dye is removed by centrifugation at 3400 × g for 3 min and cells washed once in FACS buffer before resuspending for analysis. Cells were run on a BD LSR II flow cytometer (BD Biosciences) and the results analysed using FlowJo 10 (FlowJo LLC).

### Transepithelial Electrical Resistance (TEER)

TEER values were measured using an EVOM2 resistance meter and EndOhm chamber with a 6 mm culture cup for 24-well Transwells or a 1.5 mm electrode (STX100) designed for 96-HTS Transwells (all from World Precision Instruments). Transwells were placed into the culture cup and readings were taken after the TEER reading had stabilized (typically 5-10 seconds). Readings were taken once per week for up to 4 weeks post ALI.

### Automated, High-Content Screening for Drug Efficacy and Toxicity Using HTS Transwell®-96 Permeable Supports

Recombinant GFP tagged RSV A2 strain was kindly provided by Fix et al [24] and propagated using HEp-2 cells for 3-5 days in Opti-MEM. Virus was purified as described previously [25], collected in BEBM (Life Technologies) and frozen at −80°C. For ALI culture infection, the apical surface of the ALI cultures was rinsed with medium (BEBM) and 50µL viral inoculum (MOI:1) in BEBM was applied to the apical surface for 1h at 37°C and then removed. Following infection, cells were fed basolaterally with media containing different concentrations of drugs that inhibit different stages of the RSV replication cycle: an inhibitor of RSV Fusion (F) protein (CPD23, compound 23 from [26]) that blocks viral entry to the cell, and a nucleoside inhibitor (ALS-8112, MedChemExpress) that causes lethal virus mutagenesis or disturbance of viral RNA synthesis to inhibit viral replication [27]. ALS-8112 was initially dissolved in DMSO and made up to a working concentration range of 500 to 4000 nM in Airway Epithelial Cell Growth Medium (Promocell, Heidelberg, Germany); 153 ± 76 nM was previously shown to result in a 50% inhibition (EC_50_) in RNA replication of RSV A2 in HEp-2 cells [27]. CPD23 was used at 5 to 80 nM (expected EC50 1.59 nM (personal communication). Cells were monitored daily over a 7-day period using an environmental chamber (5% CO_2_, 37 °C) connected to an inverted microscope system (Nikon Ti-E) with a 20x objective. Image acquisition was automated using NIS-Elements JOBS (Nikon) module (see **Supplementary Material 1&2**) to measure GFP+ fluorescence (as an indicator of viral replication) and fast time-lapse (100 fps) recording (for ciliary beat frequency calculation). Ciliary beat frequency is a sensitive indicator of cell toxicity [28–30] and concurrent image acquisition with Nikon NIS-Elements JOBS module allowed us to analyze drug efficacy and ciliary activity in parallel in 96-Transwell ALI cultures.

### Data Analysis and Statistical Methods

CBF was determined from timelapse files by first extracting the average pixel intensities using ImageJ (NIH) and, secondly by performing a fast Fourier transformation (FFT) on this data using R; 6400 regions of interest (ROI), each with an area of 16.8*µ*m^2^ were analysed per file (pixel resolution = 0.32µm). The ImageJ macro and R code (`ciliR` package [31]) used in this study are included in the **Supplementary Material 3**. Analysis of RSV-GFP fluorescence was performed by using a custom ImageJ macro which counted the number of infected cells based on binary thresholding (macro included as **Supplementary Material**). Statistical analyses were performed in R version 4.0.2 [32] and GraphPad Prism version 9.0 using the statistical tests indicated in figure legends. Drug combination was analysed using the `SynergyFinder version 3.14.0` package for R [33]. The Loewe model was applied to determine synergy score. The degree of smoothing in probability density estimates was determined via cross-validation.

## Results

### Evaluation and Quality Control of the Air-Liquid Interface Primary Differentiated 96-HTS Transwell Model

To evaluate the suitability of the 96-HTS ALI model for anti-viral testing compared to traditional 24-Transwell systems, we compared the cell type populations (using flow cytometry) and ciliary beat frequency (using high-speed video microscopy) of primary cells obtained from two adult individuals (n=2) grown using 96-HTS format to more traditional 24-Transwell format. Immunofluorescence imaging showed formation of a confluent epithelium containing equivalent levels of ciliated cells (β-tubulin, **Figure 2A**). Flow cytometry supported this observation where we found the proportion of basal cells and ciliated cells (% cell type from total live cells) was not significantly different between the 96-HTS and 24-well plate format at 7% and 9%, and 42% and 48.3% respectively (**Figure 2B&C**). The average (mean±SEM) ciliary beat frequency (CBF) of ciliated cells grown on 96-HTS Transwells was slightly higher (but within 1Hz margin of error) compared to the same donors’ cells grown using the 24-Transwell format (i.e. 12.43±1.99 Hz compared to 11.62±1.26Hz, respectively for one donor and 11.91±1.13 Hz and 10.68± 1.38 for the second donor). The variability of the mean CBF between all wells across the 96-HTS plate was low, with a mean (±SEM) of 11.41 ± 0.08 Hz (**Figure 2D**).

**Figure 2:**
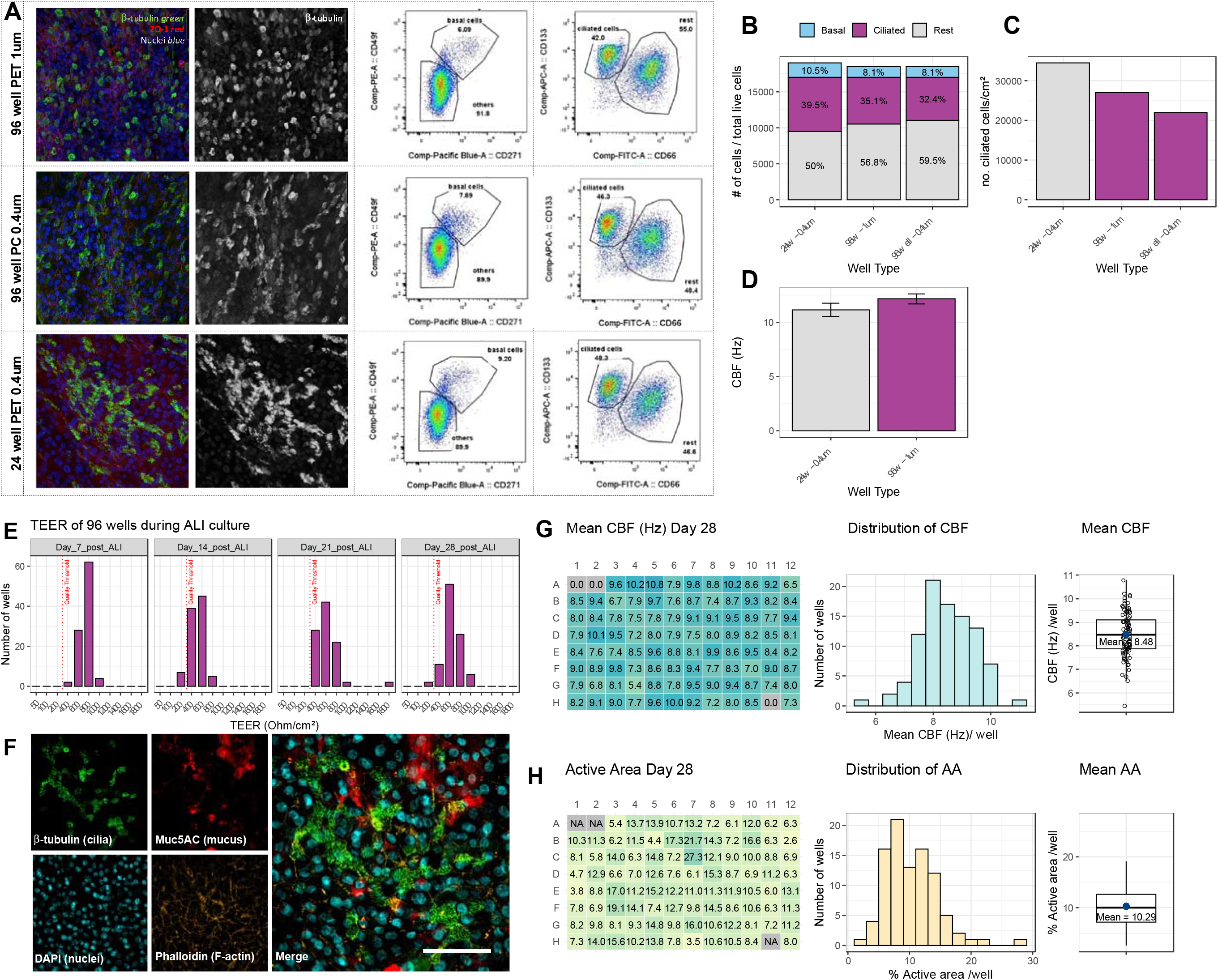
Characterization of the 96-HTS HTS ALI-culture model. (**A**) Comparison of 24-well and 96-HTS Transwell systems. Left panels show immunofluorescence images showing the presence of β-tubulin-expressing cilia (green) in 96-HTS format ALI cultures. Phalloidin (orange) stains F-actin and DAPI (blue) stains nuclei. These images are representative of two healthy donors (n=3 images). Right panels show flow cytometric evaluation of cell populations recovered from cultures grown at ALI on 96- (n=4 wells) and 24-well plates (n=2 wells) and quantified in (B) and (C). (**D**) The mean ciliary beat frequency (CBF) of cultures grown at ALI on 96- (n=2 donors) and 24-well plates (n=2 donors) as determined using the `ciliR` package. (**E**) Histograms of the trans-epithelial electrical resistance (TEER) of all wells in a 96-HTS plate of ALI culture across the 4 weeks needed for differentiation, n=96. (**F**) Representative immunofluorescence images showing the presence of β-tubulin-expressing cilia (green) and MUC5AC-expressing mucosecretory cells (red) in 96-HTS plate format ALI cultures. Phalloidin (orange) stains F-actin and DAPI (blue) stains nuclei. These images are representative of two healthy donors (n=3 images). (**G**) Example 96 well plate layout showing mean CBF per well (left panel), a histogram showing the frequency of CBF (centre panel) and mean CBF across a plate of a 96well HTS ALI cultures as determined using the `ciliR` package. (**H**) Example 96 well plate layout showing mean active area per well (left panel), a histogram showing the frequency of active area (centre panel) and mean active area across a 96well HTS ALI culture plate as determined using the `ciliR` package.

To assess the uniformity of the 96-HTS format, we measured the transepithelial electrical resistance (TEER) as the cells differentiated. We found that on Day 7 post ALI 97% of wells grown using the 96-HTS format had TEER of >300 Ωcm^2^ (median = 600 Ωcm^2^), which is regarded as sufficient for effective barrier function in quality control tests (**Figure 2E**). At day 14 and 21 post ALI 100% of wells had TEER reading >300 Ωcm^2^ (median = 800 Ωcm^2^ and 600 Ωcm^2^, respectively). At day 28 post-ALI 98% of wells had TEER reading >300 Ωcm^2^ (median = 600 Ωcm^2^) (**Figure 2E**). Immunofluorescence imaging showed formation of a confluent epithelium containing both mucosecretory cells (MUC5AC+) and ciliated cells (B-tubulin (**Figure 2F and Supplementary Figure 1**).

We utilized automated well scanning to capture fast time-lapse videos of each well in the 96-HTS plate and developed R code to assess the CBF and active area, visualizing this data as a plate map (**Figure 2G & H**, respectively). This method serves as a further quality control measure to identify and eliminate suboptimal wells, while also establishing the standard deviation in CBF across the plate. We also plotted the ciliary beating in the layout of 96 well plate (**Supplementary Figure 2**) which over repeated experiments could be used to determine experimental design faults such as plate edge effects or handling errors. This baseline variation is crucial for accurately determining the effects of drugs or viruses in subsequent experiments.

These experiments confirm that miniaturized 96-HTS ALI cultures maintain comparable epithelial integrity, cellular composition, and ciliary beat frequency to larger well formats.

#### Assessment of Drug Efficacy and Ciliary Beat Frequency Under Antiviral Drug Treatment

Next, we evaluated the 96-HTS ALI model using a recombinant RSV with a GFP reporter linked to the L protein (polymerase) [24] to provide sensitive readouts of drug efficacy by identifying infected cells in differentiated primary airway epithelial cells. This approach is novel in its potential to quantify the numbers of infected cells and evaluate drug combinations and possible synergy using primary differentiated cells in a high-throughput system. To do so, we investigated the effect of potential anti-viral drugs that inhibit different stages of the RSV replication cycle: an inhibitor of RSV Fusion (F) protein (CPD23) that blocks viral entry to the cell, and a nucleoside inhibitor (ALS-8112) that interferes with viral replication [27]. It is possible that these two mechanisms of action could act in a synergistic or additive way to enhance antiviral activity.

Image scans of whole wells were collected three days after RSV infection (**Figure 3A**), and GFP fluorescence was measured to indicate the number of infected cells. As monotherapies, the drugs showed an efficacy range of 57-98% inhibition of viral replication (n=1 donor, 3×3 replicates). The IC50 of ALS-8112 averaged 475 nM, aligning with literature values of 15-500 nM in similar ALI culture assays [34] and 20 nM in replicon assays[35]. The IC50 of CPD23 at 2.39 nM was found to be lower than the lowest concentration tested (5nM) so is reported with low confidence (**Figure 3B**).

**Figure 3.**
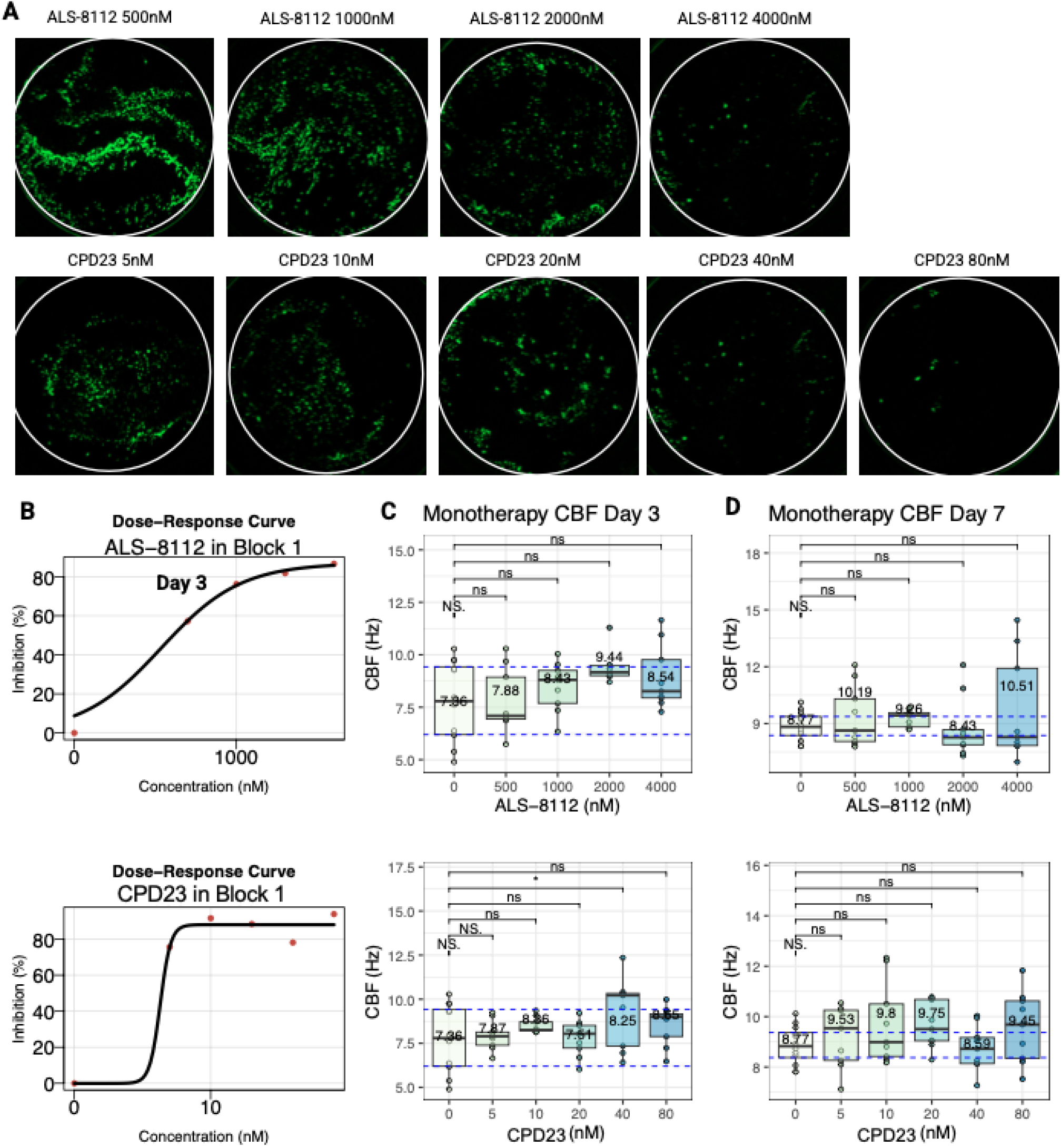
Viral Inhibition and Ciliary Beat Frequency Analysis with High Dose Monotherapies of ALS-8112 and CPD23. (**A**) Representative fluorescence images of whole wells showing GFP expression, indicating RSV infection, at various concentrations of ALS-8112 (500-4000 nM) and CPD23 (5-80 nM). Wells treated with increasing concentrations of ALS-8112 and CPD23 exhibit reduced GFP expression, indicating inhibition of RSV replication. Scale bar represents 1mm. (**B**) Dose-response curves showing inhibition of RSV replication (n=3 plates). (**C&D**) Boxplots showing the mean and distribution of CBF for ciliated cells treated with varying concentrations of ALS-8112 (500-4000 nM) (C) or CPD23 (5-80 nM) (D) 7 days post-infection. The CBF distribution remains relatively consistent across the different concentrations, suggesting minimal impact on ciliary function

Focusing on cell toxicity, we found that high concentrations of ALS-8112 did not affect (p>0.05) the ciliary beat frequency (CBF), with a mean±SD CBF of 7.36±4.7 Hz at 0 nM and 8.54±5.09 Hz at 4000 nM at Day 3 (**Figure 3C**) and 8.77±5.08 Hz at 0 nM and 10.5±5.37 at 4000 nM (± 2.1 Hz) at Day 7(**Figure 3D**). Treatment with CPD23, also did not affect the CBF, with 8.64±5.11 Hz at the highest concentration of 80 nM on Day 3 (**Figure 3C**) and 9.45±5.43 Hz at Day 7 (**Figure 3D**).

### Evaluation of Combination Anti-Viral Therapy in 96-HTS: Drug Synergy and Toxicity Analysis

An advantage of the 96-HTS ALI model is the potential to test multiple drugs or drug combinations, allowing the evaluation of the combined anti-viral effect of ALS-8112 and CPD23 over a range of doses. We generated a dose–response matrix where the response is represented as the percentage of inhibition compared to the untreated control (n=3 replicates per plate completed in triplicate 3 plates) (**Figure 4A**).

**Figure 4.**
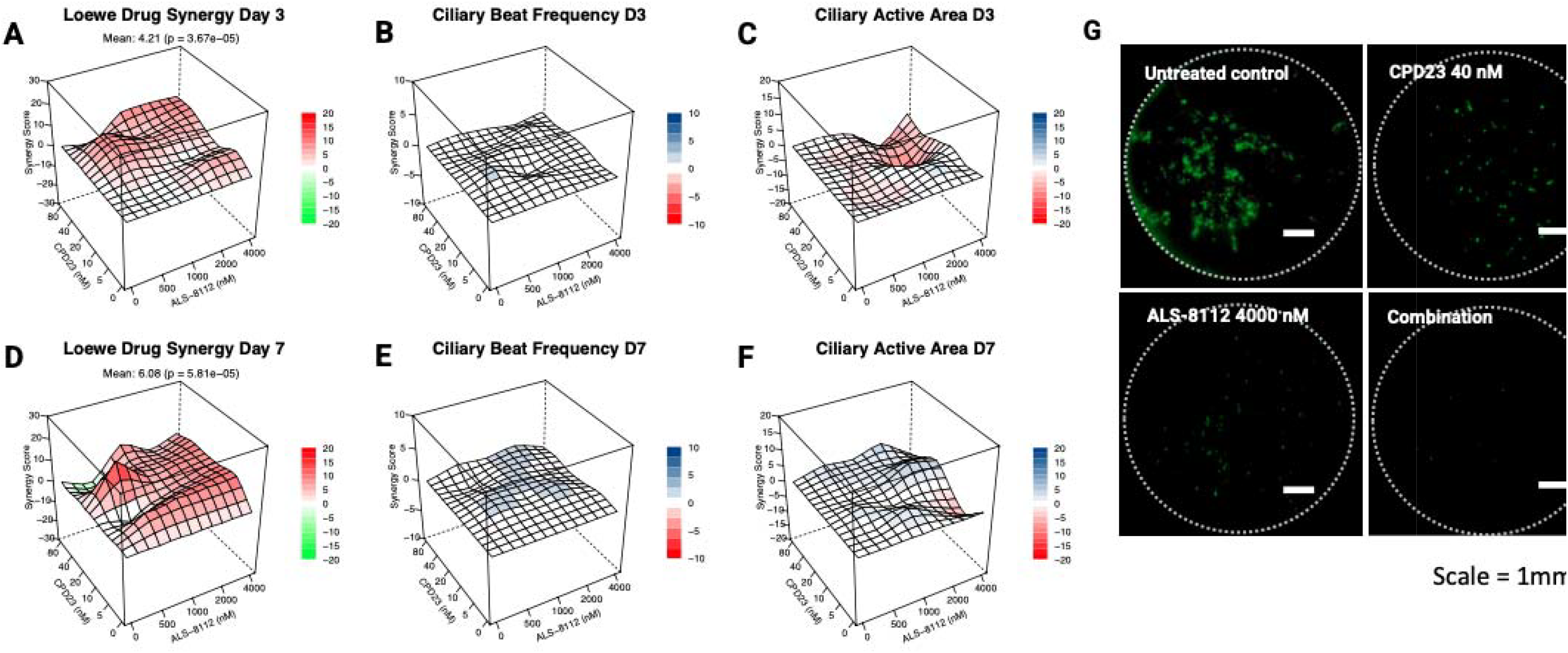
Dose-response and toxicity analysis of combination therapies in primary human airway epithelial cells. (**A**) Synergy Analysis at day 3 (**A**) and day 7 (**D**): - The 3D synergy maps illustrate the interaction between ALS-8112 and CPD23. The synergy scores were calculated using the Loewe model, where red regions indicate synergistic interactions, and green regions indicate antagonistic interactions. Overall values between −10 and 10 are considered additive. **(B&E)** Toxicity Matrices of the ciliary beat frequency (CBF) across the 96-HTS treatment plate at day 3 (B) and day 7 (E), shown as a three-dimensional surface plot. The mean CBF remained between 8-9 Hz, indicating minimal impact on ciliary activity. (**C&F**) The active area percentage, indicating overall cellular activity, remained consistent at day 3 (C) and day 7 (F) with untreated plates across different concentrations of ALS-8112 and CPD23, demonstrating no significant toxicity. **G**) Representative fluorescence images of whole wells showing GFP expression, indicating RSV infection, with ALS-8112 (4000nM) and CPD23 (40 nM) and in combination. Wells treated with ALS-8112 and CPD23 exhibit reduced GFP expression, indicating inhibition of RSV replication. Scale bar represents 1mm

Analysis of drug combination using the `SynergyFinder` package for R and Loewe model indicated an average synergy score of 4.21 and 6.08 for ALS-8112 and CPD23 at day 3 and day 7 post-infection, respectively (**Figure 4B and C**). This software applies reference models [33, 36] for synergy scores calculation that can be interpreted as the average excess response due to drug interactions beyond assumption. Values below −10 indicate the effect is antagonistic, between −10 and 10 is considered additive and above 10 the drugs effect is likely to be synergistic [33]. The 3D synergy maps (**Figure 4B&C** and **Supplementary Figure 3**) highlight the synergistic and antagonistic dose regions with a gradient of red and green, respectively. Synergy maps for the Bliss, ZIP and HAS models are shown in **Supplementary Figure 3**). This facilitates a directed comparison of the interaction between the two drugs allowing to identify the areas (combinations) that have the most synergistic effect.

We found that when used in combination, ALS-8112 and CPD23 showed an additive, but not synergistic average anti-viral effect when used in a range of 0-4000nM (ALS-8112) and 0-80nM (CPD23). The maximum additive effect (local maximum in 3D maps) was found at 500 nM for ALS-8112 and 40 nM for CPD23, both at day 3 and day 7 post RSV infection. The least effective combination (local minimum) was when ALS-8112 was used at a concentration of 500 nM and CPD23 at 10nM. However, the available assay window to observe additional effects beyond those of the single compound alone is very small and possibly within assay variability. To accurately detect any synergistic combination effects, combinations should be tested at concentrations spanning the IC50 of each compound. These findings demonstrate the proof of principle for the system.

We also evaluated the effect of the combination therapy on ciliary activity, as an indication of cellular toxicity. To visualize this, we plotted a similar 3D graph (**Figure 4B**) to that constructed by the `SynergyFinder` R package, which may allow a direct comparison between the mean ciliary beat activity and anti-viral activity of the treatment wells. This figure shows that the mean CBF across the 96 well treatment plate remained between 11-12Hz (**Figure 4B&E**). Overall, the mean CBF across the 96 well treatment plate remained consistent with a mean of 8.57 Hz at Day 3 and 8.92 at Day 7 (**Supplementary Figure 4**).

## Discussion

This study has demonstrated the effectiveness of novel analysis packages in delivering data on drug efficacy and cellular toxicity using primary ciliated airway epithelial cells with conventional inverted microscopes. These packages were used to evaluate the efficacy of combination therapy against RSV in a high-throughput, disease-relevant, human cell-based *in vitro* system. Despite primary cells being the gold standard, their use comes with disadvantages, such as cost, limited lifespan, and variability between donors, passage, or experiments. Therefore, maximizing the number of test conditions achievable with each small batch of cells is essential. We cultured primary epithelial cells at air-liquid interface using a commercially available, lower-cost (per experimental unit), 96-HTS Transwell system. This miniaturized ALI culture system resulted in comparable epithelial cell composition, with similar numbers of total live cells, progenitor and differentiated cell populations compared to cultures grown using the conventional 24-Transwell format. As we previously reported[21], there was no difference in the number of motile cilia and average ciliary beat frequency between the two culture formats.

Combining this primary cell model with an RSV-GFP reporter system allowed us to test antiviral drugs individually and in combination against RSV. We used two drugs that inhibit different stages of the RSV replication cycle: an inhibitor of RSV Fusion (F) protein (CPD23, [26]) that blocks viral entry to the cell, and a nucleoside inhibitor (ALS-8112) that causes lethal virus mutagenesis or disturbance of viral RNA synthesis to inhibit viral replication [27]. We found that ALS-8112 alone had an IC50 similar to that reported previously (15-500 nM [34]). The IC_50_ of CPD23 in our system was also within the expected range and approximately 100-fold more potent than ALS-8112. This difference in potency may be indicative of drug binding or their different mechanism of action; CPD23 acts extracellularly to block viral entry to the cell, whereas ALS-8112 interferes with virus replication following viral entry. Thus, the IC_50_ of ALS-8112 maybe higher to compensate for drug uptake.

Interestingly, treatment with high dose of ALS-8112 (13-fold higher than the IC50) did not reduce the rate of ciliary beating, which is a strong indicator of cell toxicity [37]. CPD23 also did not affect CBF at highest concentration tested, which was 60-fold higher than the IC50. Capping the analysis at 13-fold of the IC50 concentration of CPD23, or 20nM, we found no significant difference in the ciliary beat frequency (10.5 Hz v 12.2 Hz). This analysis was performed using an automated widefield fluorescence microscope, with imaging and high-speed video microscopy completed for each plate in a 2 to 3-hour window. This is advantageous as it allows multiple plates to be analysed in a relatively short time.

We showed that ALS-8112 and CPD23 result in an average additive antiviral effect for combinations using a range of 0-4000 nM (ALS-8112) and 0-80 nM (CPD23) with a peak of maximum additive effect for 500 nM of ALS-8112 combined with CPD23 at 40 nM [33]. We used the Loewe model to assess the synergistic (score >10), additive or antagonistic (score <-10) effect of drug combinations respectively. A limitation of this method is that it cannot directly assess a combination effect which is higher than the achievable effect of the individual drugs; as the concentrations of the inhibitors used alone already resulted in more than a 50% reduction in viral load. Ideally, combination analyses should be performed at concentration ranges spanning each compound’s IC50. Future studies will be directed to address this limitation.

Importantly, we have demonstrated that the 96-HTS system is effective for use with primary airway cells and have developed R pipelines to streamline and enhance the accuracy of drug combination testing and analyses, paving the way for high-throughput assessment of potential therapeutic synergies.

## Conclusion

We have shown that 96-HTS air-liquid interface cultures can be used to model the airway epithelial response to RSV infection during anti-viral therapy. We anticipate that this system will be widely applicable to other respiratory viruses that show tropism for epithelial cells, such as SARS-CoV-2 and influenza A virus. In the future, larger scale combinatorial screening of drug therapies using models derived from vulnerable groups such as infants and patients with primary immunodeficiencies has the potential to rapidly inform the clinical translation of effective drug regimens targeting respiratory RNA viruses.

## Acknowledgements

This work was funded by a MedCity Collaborate to Innovate grant (awarded to C.M.S and E.T, C2N 543922). C.M.S also acknowledges funding support from UKRI/ BBSRC (BB/V006738/1). This work was supported by the NIHR Great Ormond Street Hospital Biomedical Research Centre. The views expressed are those of the author(s) and not necessarily those of the NHS, the NIHR or the Department of Health. Analysis was performed at the Light Microscopy and Flow Cytometry core facilities at UCL Great Ormond Street Institute of Child Health, which are supported by the NIHR GOSH BRC award 17DD08. The views expressed are those of the authors and not necessarily those of the NHS, the NIHR or the Department of Health.

## Author contributions

DC designed the study, conducted experiments, analysed data, and cowrote the manuscript. OG, MC-B and CMS co-designed the `ciliR` package and performed the CBF analysis. DDHL and COC provided support through ethics and donor recruitment. RA, ET and CMS conceived the study and oversaw the funding application. ET, REH and CMS oversaw data analysis and interpretation, and the write-up of the manuscript.

## Code availability

Custom code for the analysis performed in this study is publicly available via GitHub at https://github.com/smithlab-code/ciliR.

