## Supplementary figures and images for "Donor-derived airway ALI model for high-throughput screening of antiviral combinations with concurrent analysis of antiviral efficacy and epithelial toxicity using ciliR"

### Supplementary Figure 1

Supplementary Figure 1

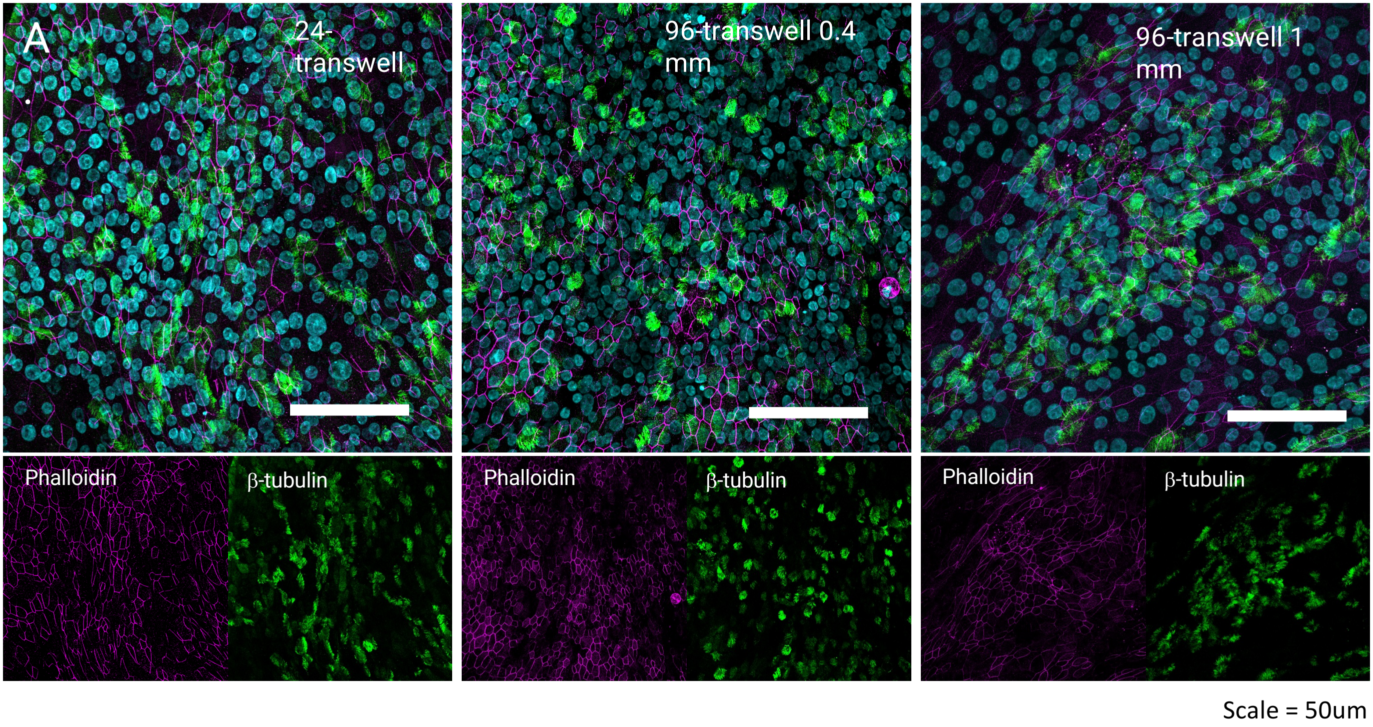

### Supplementary Figure 2

Supplementary Figure 2

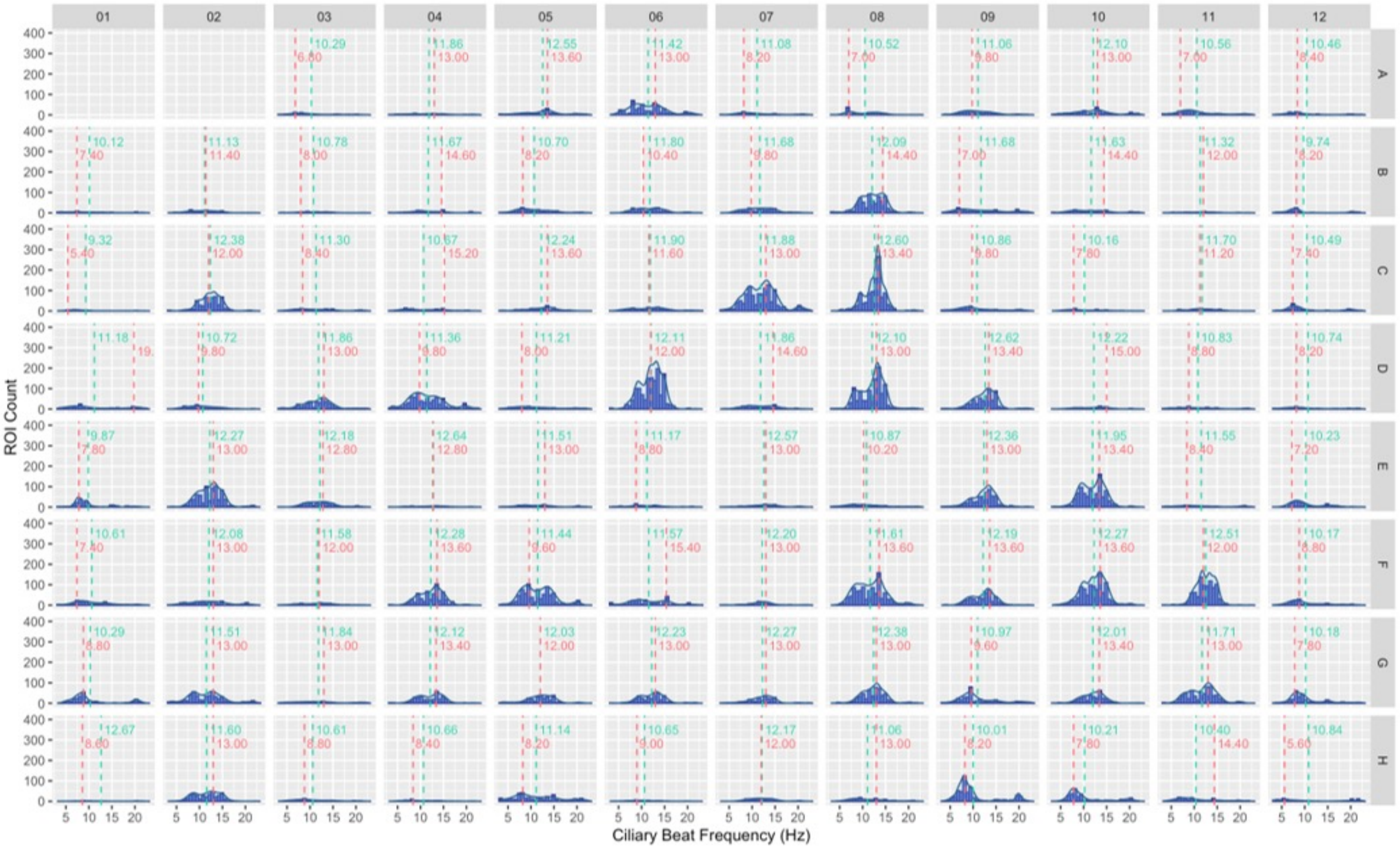

### Supplementary Figure 3

**Supplementary Figure 3**

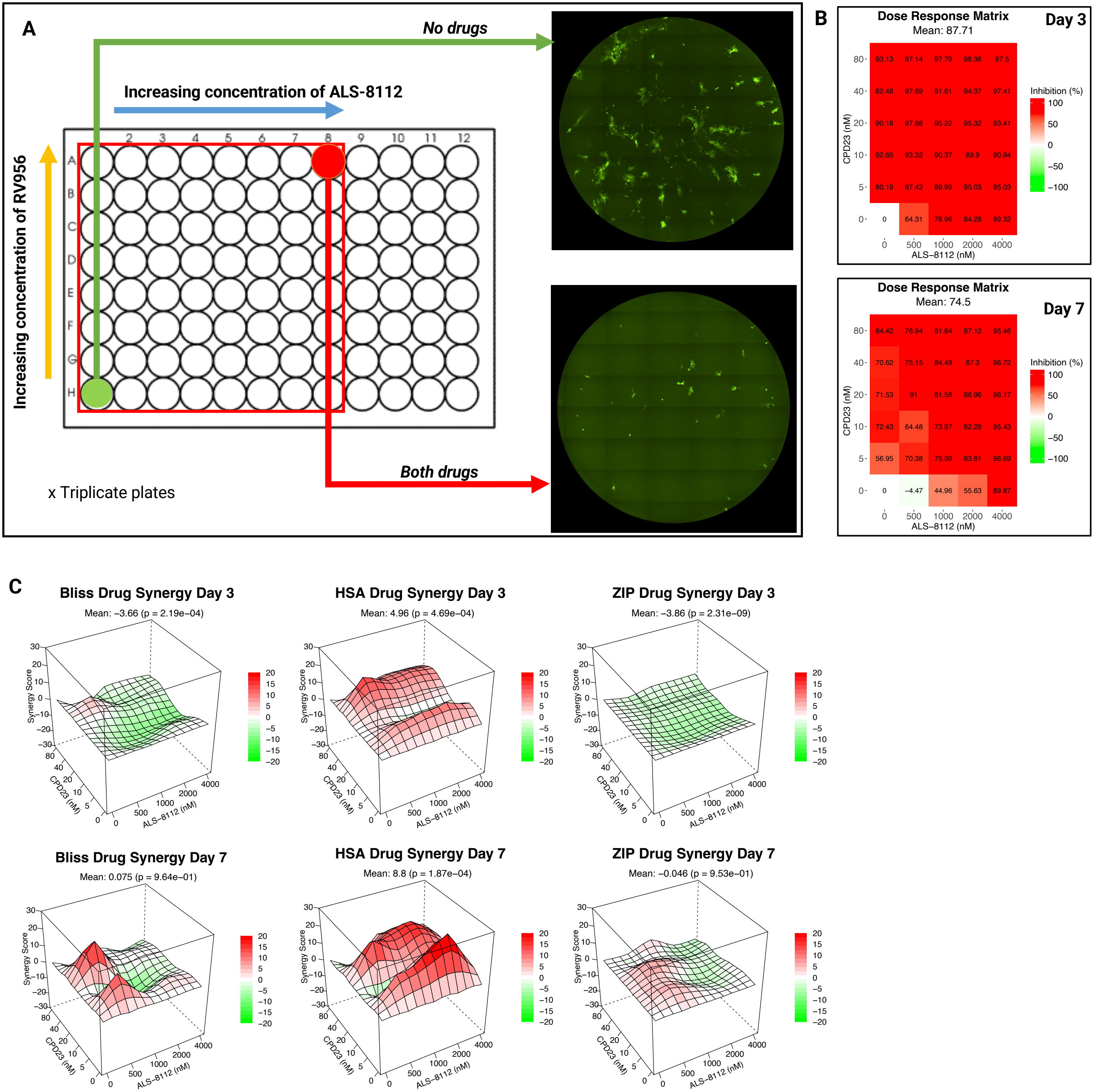

### Supplementary Figure 4

#### Supplementary Figure 4

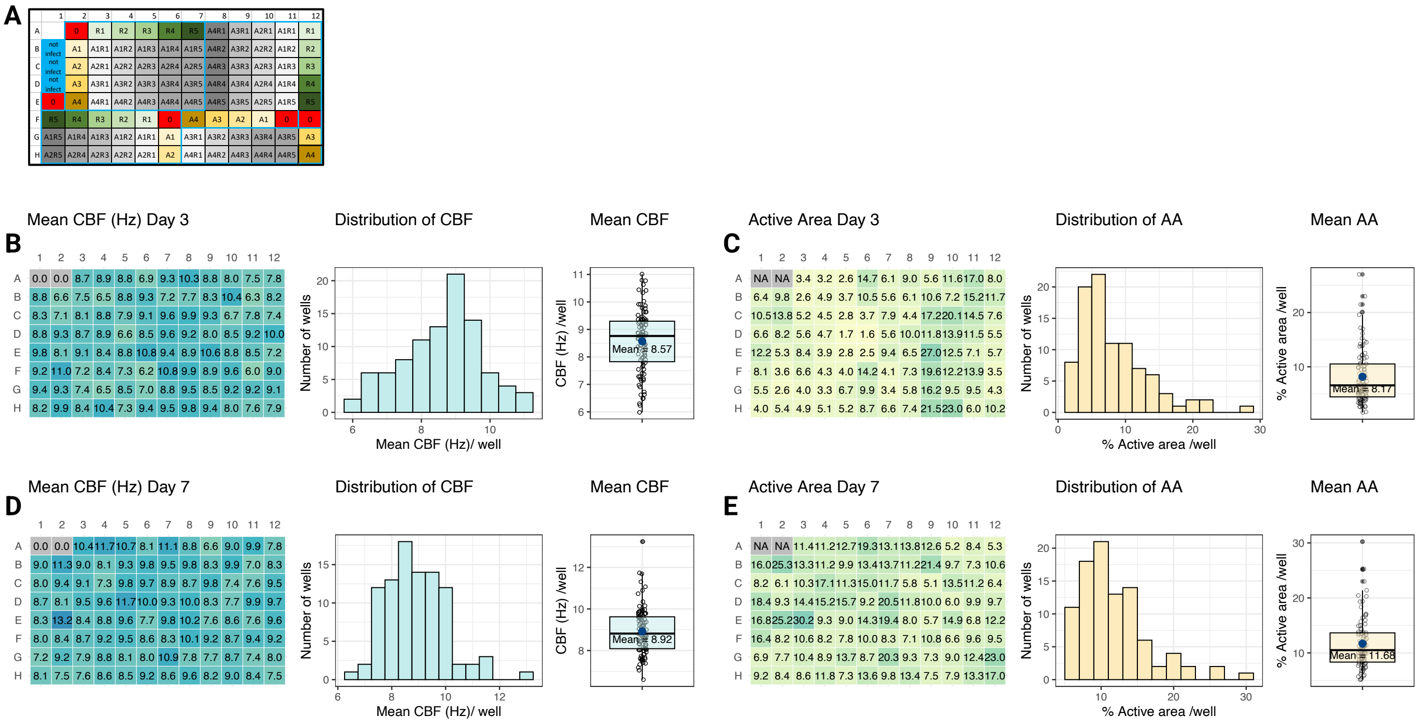
