## Supplementary Material for "Donor-derived airway ALI model for high-throughput screening of antiviral combinations with concurrent analysis of antiviral efficacy and epithelial toxicity using ciliR"

**Supplementary Material 1**. The graphical user interface (GUI) of the Nikon Elements JOBS experimental setup for automated whole well fluorescence capture. The interface is organized into three panels; The left panel handles general settings and initial conditions, including experiment identity and timing configurations. The middle panel manages plate and wells configuration, featuring a visual representation of a 96-well plate with color-coded wells. The right panel offers advanced settings and protocols, allowing customization of experimental procedures and capture acquisition of FITC channel.

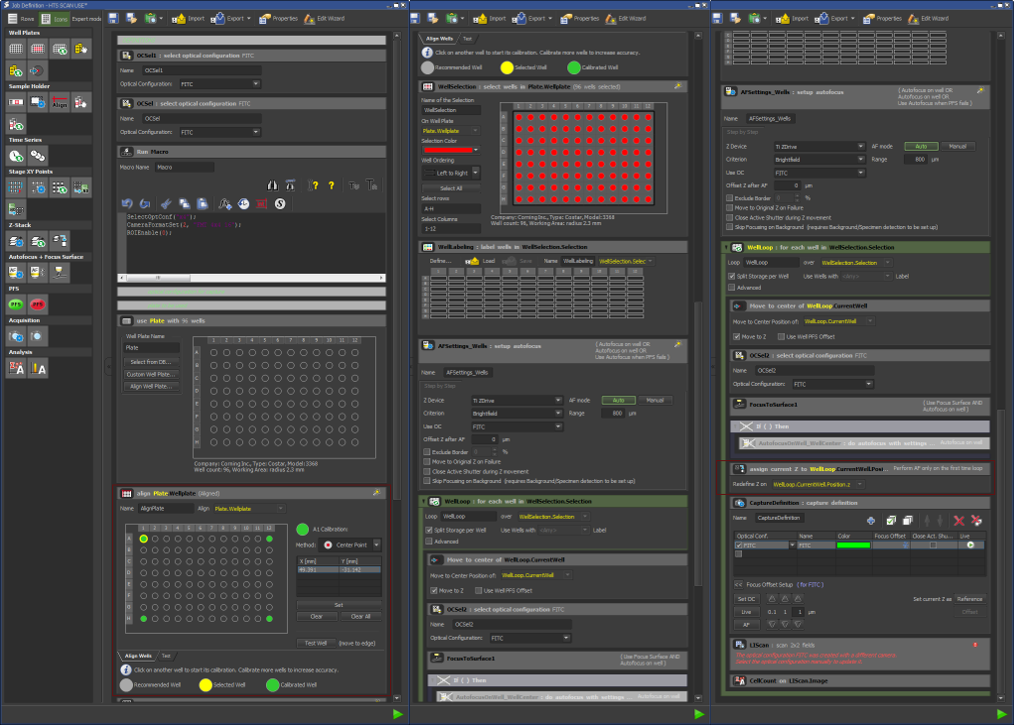

**Supplementary Material 2**. This figure shows an updated GUI for of the Nikon Elements JOBS software for automated experimental setup. Differences to Fig 1 include the addition of PFS and focus adjustment on the middle panel, and new "Points" and "Fast Timelapse" for high-speed video capture sections in the right panel.

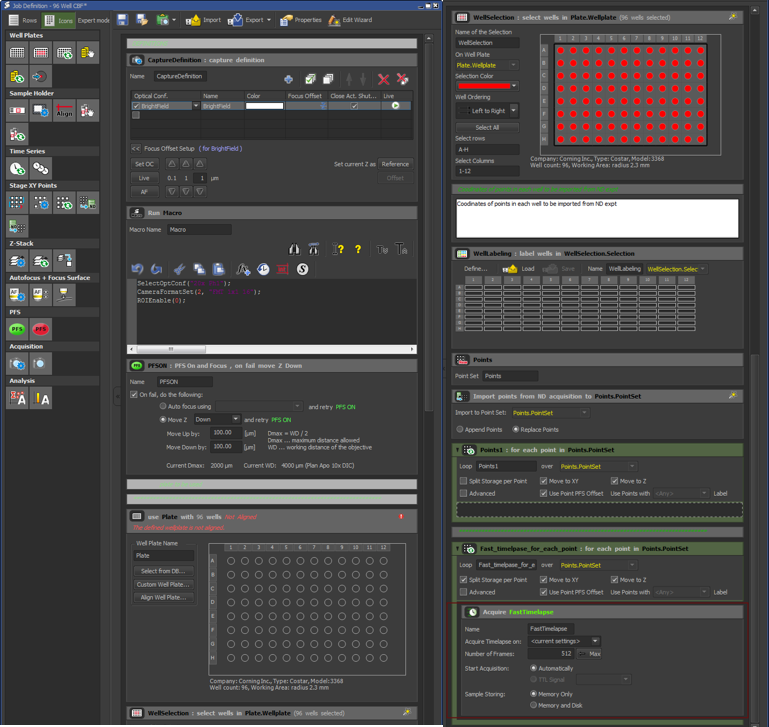

**Supplementary Material 3**

This macro for FiJI (ImageJ) automates the analysis of GFP images in a 96-well plate. It prompts the user to select a directory, processes each image by subtracting background, thresholding, and converting to a mask. It then performs watershed segmentation and particle analysis, saving intensity and count results for multiple channels (H, G, F, E, D, C, B, A). The macro creates directories for these results and saves summary files for both intensity and count data. Finally, it clears the results table and displays a completion message, streamlining the image analysis workflow and organizing the output for easy review.

```
//This macro presents a cropped and clean version of your x10 mag scan of a well of a 96 well plate. It will run through all the images in a folder.

dir1=getDirectory("Choose Image Directory");
list = getFileList(dir1);
for (i=0; i<list.length; i++)
	{
	if (File.isDirectory(dir1+list[i])){}
	else{

path = dir1+list[i];
	if (endsWith(path, ".db")){}
	else	{
		if (endsWith(path, ".ids")){}
		else	{
			run("Bio-Formats Importer", "open=[path] autoscale color_mode=Default view=Hyperstack stack_order=XYCZT");
				{

title1 = File.nameWithoutExtension;
setBatchMode(true);
dir2 = dir1+" Intensity Results"+File.separator;
File.makeDirectory(dir2);
dir3 = dir1+" Count Results"+File.separator;
File.makeDirectory(dir3);
run("Subtract Background...", "rolling=20");
saveAs("Tiff", dir2+title1+ " IntensityH.tif");
setAutoThreshold("Default dark");
run("Convert to Mask");
run("Watershed");
run("Analyze Particles...", "size=900-10000 circularity=0.30-1.00 show=Outlines display clear summarize");
saveAs("Tiff", dir3+ title1+" CountH.tif");
close();
close();

run("Subtract Background...", "rolling=20");
saveAs("Tiff", dir2+title1+ " IntensityG.tif");
setAutoThreshold("Default dark");
run("Convert to Mask");
run("Watershed");
run("Analyze Particles...", "size=900-10000 circularity=0.30-1.00 show=Outlines display clear summarize");
saveAs("Tiff", dir3+ title1+" CountG.tif");
close();
close();

run("Subtract Background...", "rolling=20");
saveAs("Tiff", dir2+title1+ " IntensityF.tif");
setAutoThreshold("Default dark");
run("Convert to Mask");
run("Watershed");
run("Analyze Particles...", "size=900-10000 circularity=0.30-1.00 show=Outlines display clear summarize");
saveAs("Tiff", dir3+ title1+" CountF.tif");
close();
close();

run("Subtract Background...", "rolling=20");
saveAs("Tiff", dir2+title1+ " IntensityE.tif");
setAutoThreshold("Default dark");
run("Convert to Mask");
run("Watershed");
run("Analyze Particles...", "size=900-10000 circularity=0.30-1.00 show=Outlines display clear summarize");
saveAs("Tiff", dir3+ title1+" CountE.tif");
close();
close();

run("Subtract Background...", "rolling=20");
saveAs("Tiff", dir2+title1+ " IntensityD.tif");
setAutoThreshold("Default dark");
run("Convert to Mask");
run("Watershed");
run("Analyze Particles...", "size=900-10000 circularity=0.30-1.00 show=Outlines display clear summarize");
saveAs("Tiff", dir3+ title1+" CountD.tif");
close();
close();

run("Subtract Background...", "rolling=20");
saveAs("Tiff", dir2+title1+ " IntensityC.tif");
setAutoThreshold("Default dark");
run("Convert to Mask");
run("Watershed");
run("Analyze Particles...", "size=900-10000 circularity=0.30-1.00 show=Outlines display clear summarize");
saveAs("Tiff", dir3+ title1+" CountC.tif");
close();
close();

run("Subtract Background...", "rolling=20");
saveAs("Tiff", dir2+title1+ " IntensityB.tif");
setAutoThreshold("Default dark");
run("Convert to Mask");
run("Watershed");
run("Analyze Particles...", "size=900-10000 circularity=0.30-1.00 show=Outlines display clear summarize");
saveAs("Tiff", dir3+ title1+" CountB.tif");
close();
close();

run("Subtract Background...", "rolling=20");
saveAs("Tiff", dir2+title1+ " IntensityA.tif");
setAutoThreshold("Default dark");
run("Convert to Mask");
run("Watershed");
run("Analyze Particles...", "size=900-10000 circularity=0.30-1.00 show=Outlines display clear summarize");
saveAs("Tiff", dir3+ title1+" CountA.tif");
close();
close();

selectWindow("Summary");
saveAs("Text", dir1+ " Count Summary.xls");

};
};
};
};
};

selectWindow("Results");
run("Clear Results");

run("Measure...", "choose=[dir2]");
selectWindow("Results");
saveAs("Text", dir1+ " Intensity Results.xls");
showMessage("ImageJ Processing Complete!")
```
